# Early reactivation of medial temporal lobe neurons during emergence from propofol anesthesia in neurosurgical patients

**DOI:** 10.64898/2026.05.22.727285

**Authors:** Joel I. Berger, Matthew I. Banks, Emily R. Dappen, Urszula J. Gorska-Klimowska, Rashmi N. Mueller, Hiroto Kawasaki, Corey A. Amlong, Melanie Boly, Wendell B. Lake, Woodrow L. Shew, Kirill V. Nourski

## Abstract

Although much is known about the molecular and cellular effects of general anesthetics, the neural mechanisms underlying loss and recovery of consciousness during anesthesia remain elusive. We provide the first report of human single neuron activity recorded throughout emergence from general anesthesia. Following cessation of propofol infusion, emergence was assessed as motor response to verbal command (RC). In most cells, firing rates increased during the course of emergence. Analysis of pooled firing rate changes indicated a shift towards criticality during emergence and attractor states during and immediately following anesthesia and immediately after RC. Changes in activity in some regions occurred prior to overt RC, with shortest latencies in the hippocampus, parahippocampal gyrus, and amygdala, followed by cingulate and then insular cortex. This work reveals that brain activity in medial temporal regions may presage restoration of responsiveness in humans following general anesthesia.

## Introduction

The neural mechanisms governing transitions between states of consciousness, for example during anesthesia induction and emergence, remain elusive. Recovery of responsiveness during emergence occurs at a lower anesthetic concentration than the loss of responsiveness during induction, indicating that emergence is not merely a result of anesthetic washout. This hysteresis, reflecting a phenomenon termed *neural inertia* [1, 2], suggests that emergence is an active and regulated process[3]. Preclinical studies have demonstrated that emergence involves transitions through a sequence of metastable states[4]. However, comparable data in human participants is limited, underscoring the need to characterize the spatiotemporal pattern of neuronal activity that mediates recovery of consciousness and responsiveness following anesthesia.

Determining the specific spatiotemporal sequence of neuronal reactivation during emergence is challenging. Scalp electroencephalography (EEG) lacks spatial precision, especially for midline and deep structures critical for consciousness and sense of self, the latter denoting one’s perception of being a distinct and autonomous agent with an identity that persists through time. Functional magnetic resonance imaging (fMRI), though spatially precise, is difficult to apply during anesthesia emergence. Intracranial recordings with hybrid micro–macro electrodes uniquely enable monitoring of single-neuron activity in these regions. Single neuron recordings provide the highest spatiotemporal resolution available *in vivo* in humans and capture heterogeneity both within and between recorded regions.

Loss of consciousness during general anesthesia is not a unitary state but a constellation of behavioral endpoints, including sedation, amnesia, hypnosis, antinociception, and paralysis, each with distinct neural substrates and dose dependencies. These selective effects are reflected in variable effects of general anesthesia on components of consciousness and sense of self. For example, consciousness as defined by the presence of subjective experience can occur in the absence of connection to the environment (i.e., dreaming), volition, or memory formation, and can be accompanied by depersonalization and disrupted sense of body ownership (i.e., out of body experiences)[5–7]. Specific aspects of consciousness and sense of self have been linked to specific brain regions, e.g., hippocampus for memory[8], amygdala for emotional salience, and insula and posterior midline cortical structures for interoceptive awareness and the sense of bodily self[9, 10].

Region- and function-specific effects of anesthetics suggest that as patients transition into and out of anesthesia-induced unconsciousness, they may pass through intermediate states in which particular aspects of consciousness and self are selectively preserved. Indeed, it has been proposed that a “core self”, i.e., the fundamental sense of existence as an autonomous entity, may persist even in the absence of consciousness under moderate anesthesia[5]. Thus, careful study of brain activity during transitions into and out of general anesthesia may yield insights into the hierarchical structure of consciousness and self. Models of self are typically hierarchical, progressing from a core sense of existence through embodied selfhood and agency to narrative identity and autobiographical memory[5, 11]. This framework raises the question: does the return to wakeful consciousness reflect a sequential reactivation of brain regions mirroring the hierarchical reassembly of the self?

Emergence from anesthesia is also likely to manifest as changes in the distance of brain networks from criticality, i.e., neural population dynamics near a critical point characterized by scale-invariant spatial and temporal correlations. The multiscale correlations conferred by criticality may support the emergence of coordination across brain regions required for the reassembly of the sense of self. A system near criticality, the boundary between order and disorder, maintains a balance between integration and segregation, maximizing dynamic range and information transmission[12, 13]. Signatures of near-criticality have been reported in the awake brain (e.g., [14]), and deviations from criticality are observed during altered states of consciousness including anesthesia-induced unresponsiveness[15–17]. Here, we quantified proximity to criticality using a recently developed temporal renormalization group approach that estimates *d*_2_, an information-theoretic distance from the nearest critical manifold, from multi-neuron spike count time series[18].

These changes in proximity to criticality unfold within a broader dynamical landscape in which brain networks progress between stable attractor states along trajectories through a high-dimensional state space[19–21]. This is particularly relevant to emergence from anesthesia, where ‘neural inertia’, i.e., the tendency of the brain to resist state change beyond what is expected from pharmacokinetics, has been observed[1, 2]. This is in contrast to induction of anesthesia, which is more predictable and occurs rapidly; emergence exhibits substantial variability across individuals, both in duration and in postanesthetic cognitive outcomes[22]. The variable time course and cognitive outcomes may depend on the specific sequence of network states traversed between the time anesthesia is turned off and patients first become responsive[4, 23]. Evidence from preclinical models suggests that emergence proceeds through abrupt transitions between discrete metastable network states (i.e. attractor states) rather than a smooth, continuous return to wakefulness [4]. Whether analogous attractor dynamics operate at the level of individual neurons in the human brain during emergence has not been examined.

Our previous work has shown that the intravenous anesthetic propofol globally disrupts differentiation and integration in the brain[24]. Propofol also specifically modulates activity and connectivity in brain networks associated with consciousness and arousal, including the default mode, executive control, and salience networks[25–27]. Here, we focus on the dynamics of neuronal activity during emergence from anesthesia. We examined the activity of single neurons in medial temporal lobe (MTL), prefrontal, and parietal cortex while participants transitioned from propofol anesthesia to an awake, responsive state. We hypothesized that brain activity during emergence from anesthesia would transition from a state that is further from criticality to one that is close to criticality when patients exhibited overt responsiveness, following a distinct trajectory of neural activity over time. Further, consistent with hierarchical models of the sense of self and self-related processing (e.g.[5, 11]), we expected return of responsiveness would rely on early re-establishment of neural activity in brain regions underlying core aspects of embodied self and awareness, and return of higher order aspects of identity such as autobiographical memory would occur later in time. Thus, we hypothesized that changes in neural activity in insula[10] and posterior cingulate cortex[9] would precede overt behavioral responsiveness during emergence and changes in activity in MTL structures[8]. Contrary to this prediction, we observed that neurons in MTL structures, including hippocampus and amygdala, changed their firing rates earliest during emergence, preceding changes in insula and cingulate cortex.

## Results

Neuronal spiking activity was recorded in 11 participants (**Supplementary Table 1**) during emergence from propofol anesthesia (**Figure 1a**). A total of 356 putative single neurons were resolved in several mesial cortical and subcortical regions, with largest representation in hippocampus, amygdala, insula, and cingulate cortex (**Figure 1b; Supplementary Table 1**). Single neurons identified by our spike sorting algorithm met criteria for waveform consistency over time and refractory periods (**Supplementary Figure 1**). Amplitude distributions for single neurons showed no evidence of missing small events (**Supplementary Figure 2**), indicating that the detection algorithm was robust. When single neurons were isolated on a particular microwire, the yield was between 1 and 8 single neurons, consistent with previous work [28]. Various other quality metrics for all isolated neurons are shown in **Supplementary Figure 1**.

**Figure 1.**
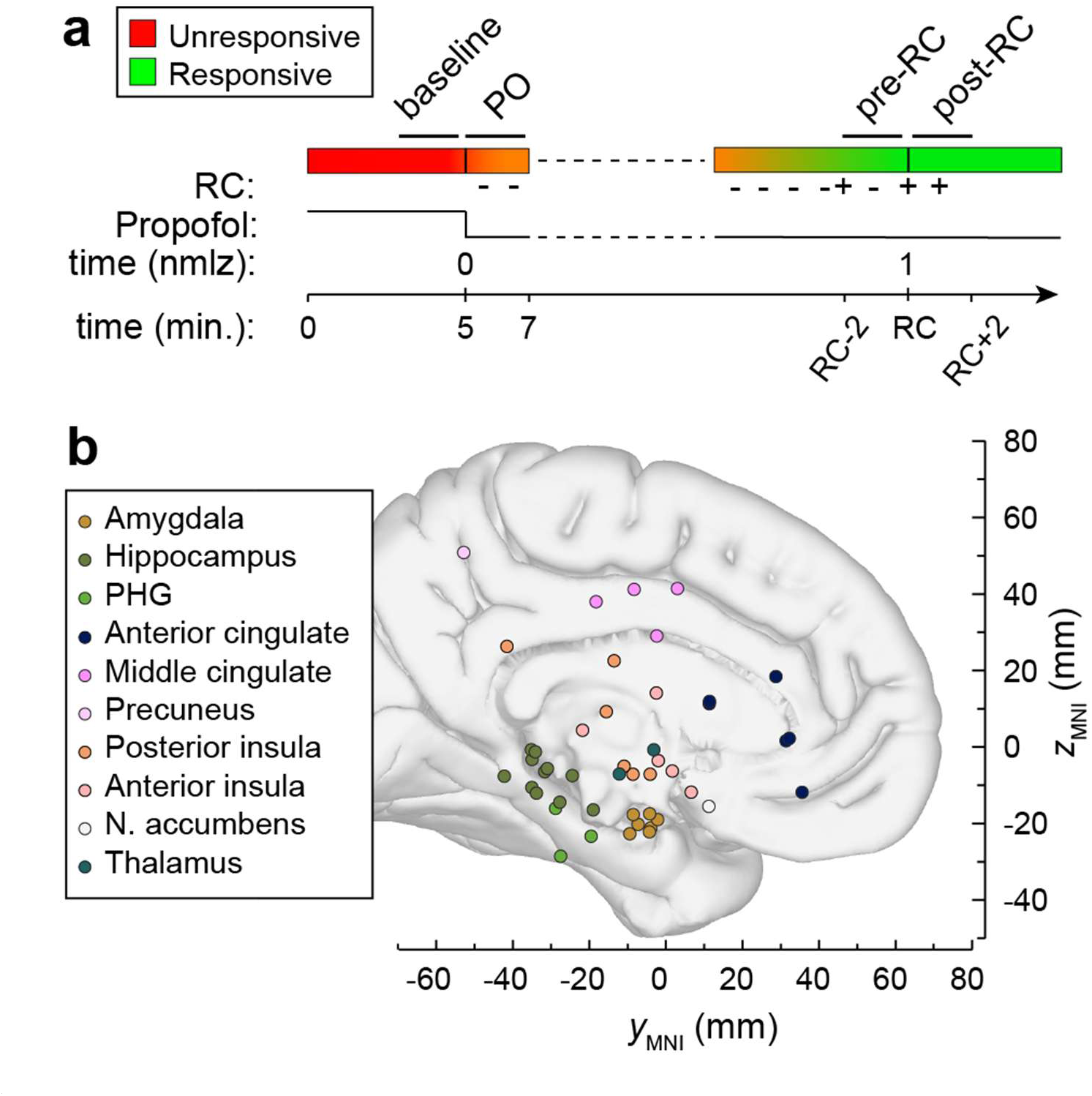
Experimental details. (**a**) Timeline of experimental protocol. From bottom: time in minutes (t_RC_: time of response to command), time normalized (nmlz) to time of propofol-off (PO) and first response to command (RC; 1), representation of propofol dose (=0 at PO), and color line depicting gradual transition from unresponsive (red) to responsive (green). (**b**) Anatomical distribution of neurons. Each filled symbol represents the location of a microwire bundle in one participant. Locations are plotted in MNI coordinate space and projected onto the left hemisphere mesial view of the FreeSurfer average template brain for spatial reference.

Behavioral responsiveness during emergence was assessed approximately every minute by asking each participant to squeeze the research team member’s hand. Return of responsiveness (i.e., ‘connected consciousness’[29]) was defined as the time of the first of two consecutive positive responses. We defined that time as response to command (RC). Total emergence time, i.e., the time between cessation of propofol infusion and RC, varied widely between individuals, ranging from 7.01 to 38.15 min. (median = 10.19 min.; interquartile interval [IQR] = 7.29). This level of inter-individual variability in time-to-emergence from propofol anesthesia is consistent with other literature (e.g.,[30, 31]).

Firing rates were calculated using a 1-second sliding window during the experiment to investigate changes in the activity of individual single neurons during emergence (**Figure 2a**). Spiking activity changed systematically during emergence, with most neurons increasing their firing rate from baseline but a substantial minority decreasing activity (see example in **Figure 2b**; also **Supplementary Figure 3**).

**Figure 2.**
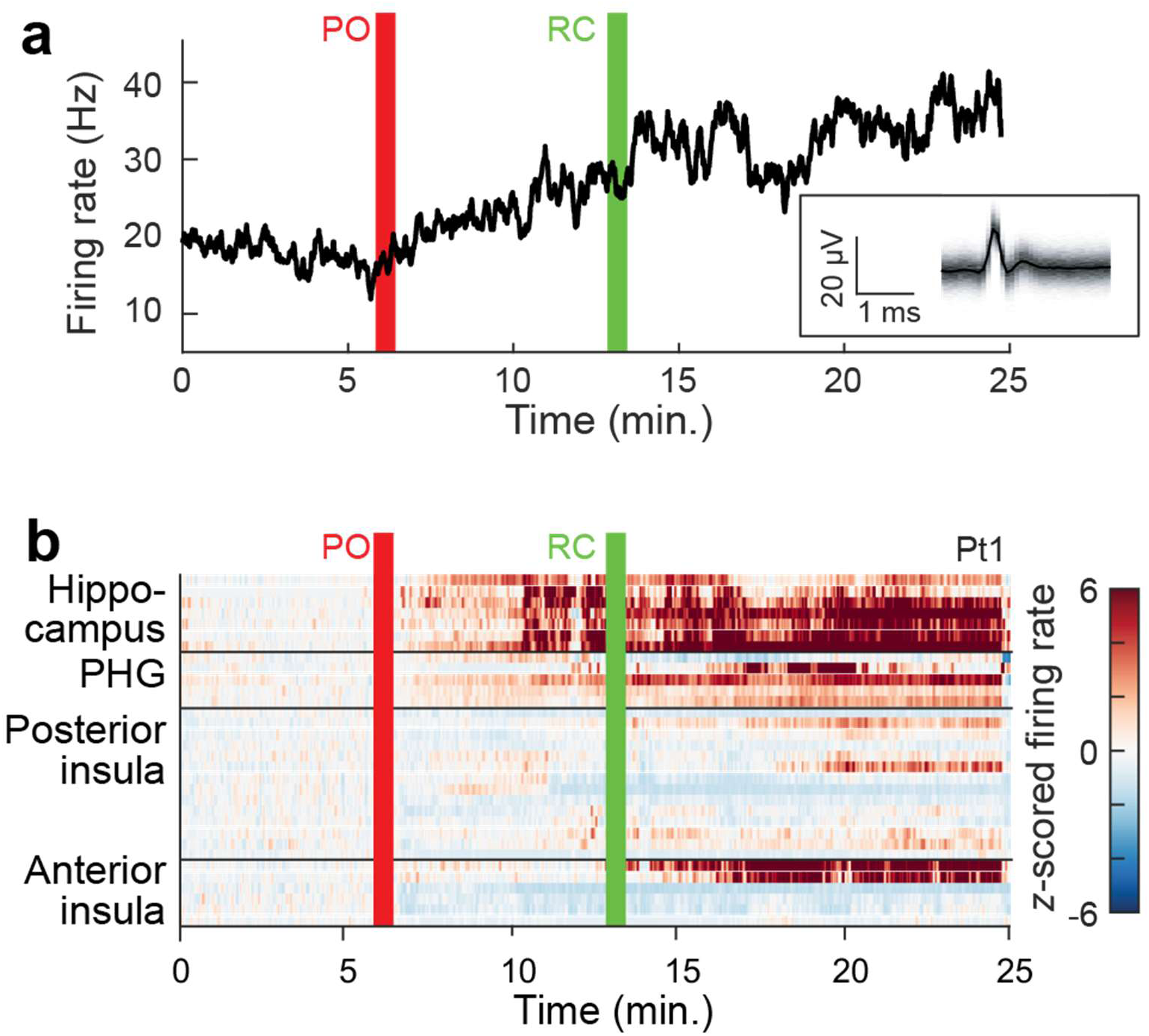
Dynamics of neuronal firing during emergence. (**a**) Example single neuron activity during emergence plotted as firing rate over time during the experiment. Firing rate was calculated in 1 s bins and smoothed with a 5 s moving average filter. *Inset* shows average spike waveform and density across all detected waveforms for that neuron. PO: propofol off. RC: response to command (i.e. the time of the first of two consecutive positive responses). (**b**) Firing rate for all neurons recorded during the same experiment in **a**. Each row corresponds to one neuron. Firing rates were *z*-scored and are shown as heat maps.

To gain insight into the collective neural dynamics underlying the transition from unresponsiveness to connected consciousness, we performed two analyses. First, we computed the distance to criticality (*d*_2_) using a recently introduced method that grounds the measurement of proximity to criticality in a rigorous theoretical framework[18]. Briefly, a vector autoregressive model is fit to multi-neuron spike count time series, and *d*_2_ is defined as an information-theoretic distance (Kullback-Leibler divergence rate) from the best fit model to the nearest critical manifold, which is determined based on temporal renormalization group theory. Smaller values of *d*_2_ indicate that population spiking dynamics are closer to criticality. For each region of interest (ROI) containing at least five isolated single neurons, spike trains were concatenated across neurons and binned at 0.2-sec resolution, and *d*_2_ was estimated in 40-sec sliding windows. By defining *t* = 0 as PO time and *t* = 1 as RC, time axis normalization enabled averaging across ROIs and participants; the pooled data showed clear evidence of decreasing distance to criticality during emergence (**Figure 3**). During deep anesthesia (*t* < 0), *d*_2_ was stable at approximately 0.25, consistent with neural dynamics substantially removed from criticality. Following return of responsiveness (*t* > 1), *d*_2_ stabilized at a lower value of approximately 0.17, reflecting dynamics closer to the critical manifold in the awake state. Between these epochs, *d*_2_ decreased across the emergence window, suggesting that the recovery of consciousness is accompanied by a gradual, continuous shift of cortical population dynamics toward criticality.

**Figure 3.**
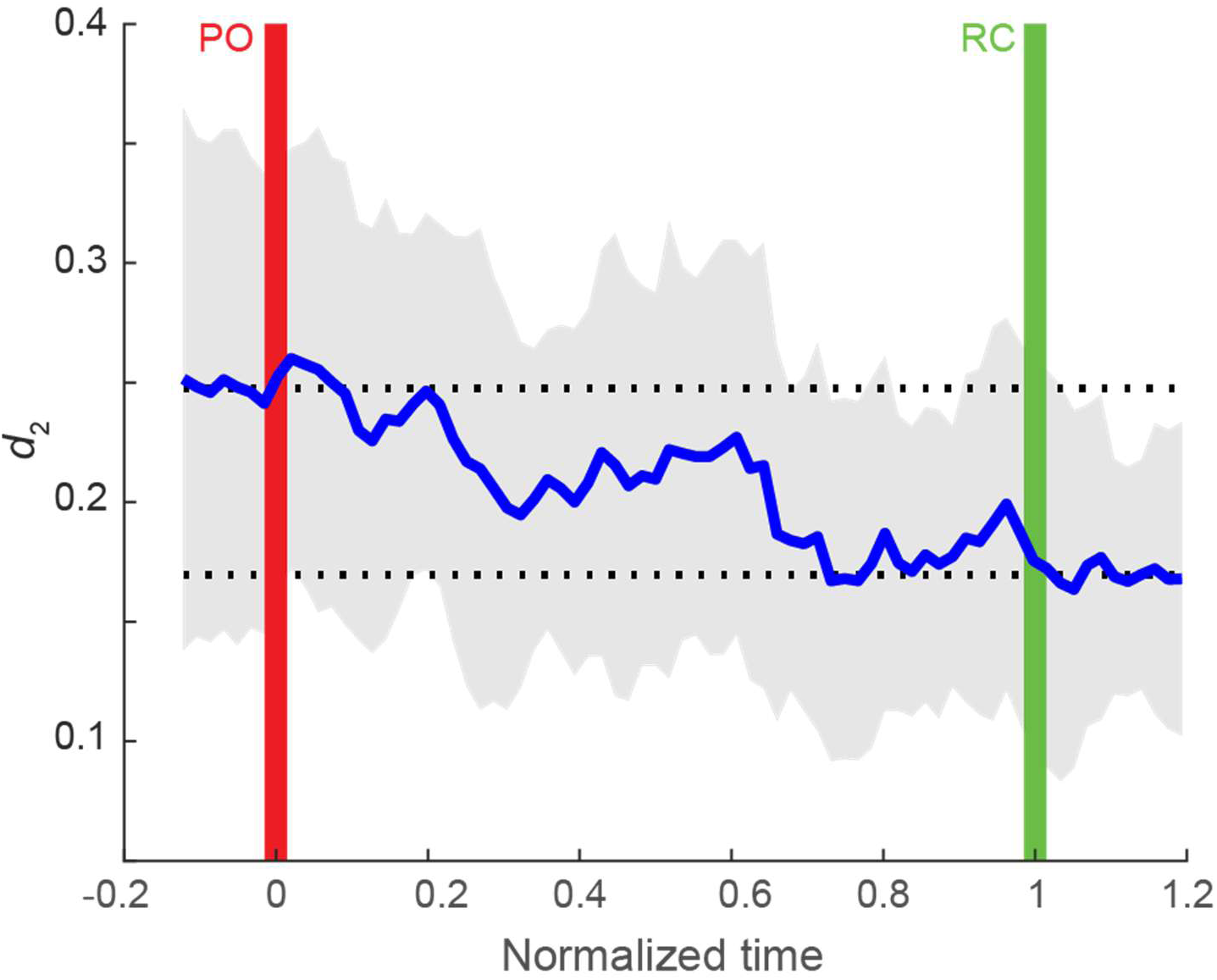
Decreased distance to criticality during emergence. *Blue line* shows *d_2_* averaged across all ROIs and all participants. *Grey shading* shows one standard deviation above and below the mean. Upper and lower *dotted horizontal lines* show mean values of *d_2_* for *t*<0 and *t*>1, respectively. PO: propofol off. RC: response to command.

In the second analysis, single neuron activity was pooled across participants by forming a time-varying firing rate vector with dimension equal to the number of recorded neurons and time resolution set by the sliding window analysis. Principal component analysis was then used to identify a 3-dimensional manifold along which neuronal activity followed a trajectory that developed over time (**Figure 4a**). Neurons from hippocampus and amygdala, the two regions with overall shortest latencies to firing rate changes, exerted the most influence on the course of the overall firing rate trajectory (**Supplementary Figure 4**). Attractor states were identified by measuring the density of the trajectory as a function of time was used to identify putative attractor states, which appear as areas of high density (**Figure 4b**). Two attractor states were identified: one while propofol was still on, and one shortly after RC, when connected consciousness was restored.

**Figure 4.**
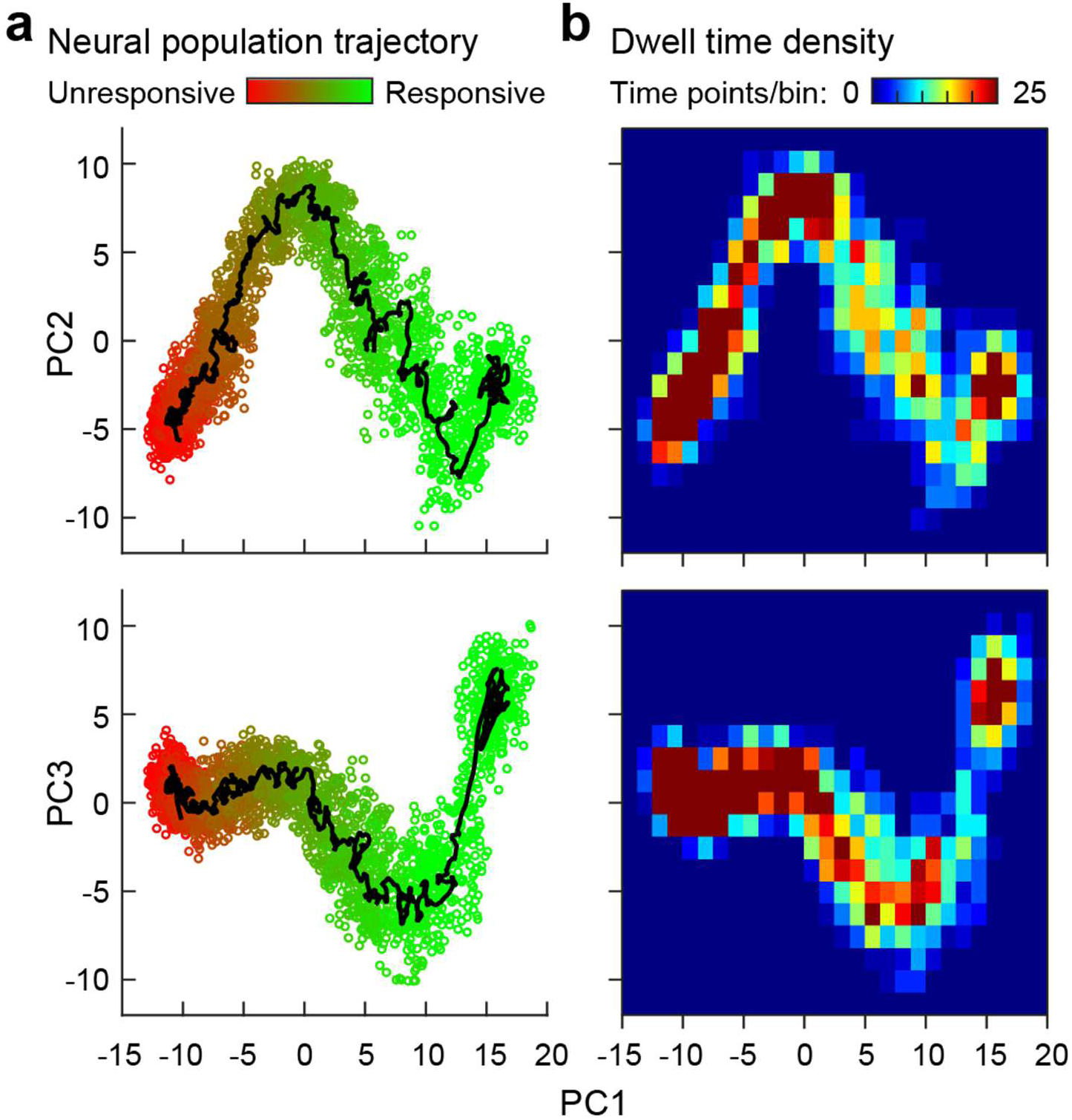
Dynamics of global neuronal activity during emergence. (**a**) State space analysis of firing rates (as in Figure 2a) across all recorded neurons, showing changes in the first three principal components (PC) as participants transition from unresponsive (*red*) to responsive (*green*). (**b**), Evidence for attractor states during emergence. Shown is the dwell-time density of the corresponding panel in **a** (see Methods). Attractor states are implicated by the presence of dark red clusters in the right subpanels.

To summarize firing rate data across regions and participants, four 2-minute time windows of interest were identified (**Figure 1a**): baseline (immediately prior to turning propofol off), propofol off (PO, immediately after turning propofol off), immediately prior to RC (pre-RC), and immediately following response to command (post-RC). Neurons were included in this analysis where they had a minimum firing rate ≥0.1 Hz in each of these windows. There was a high degree of heterogeneity in the modulation patterns across neurons and ROIs (**Figure 5**). On average, the largest increases at the post-RC timepoint were observed in hippocampus and the smallest in posterior and anterior insula. Statistical testing of these ratios based on a linear-mixed effects model with post-hoc testing (see *Methods*) showed that firing rates during the post-RC window were significantly elevated relative to baseline in hippocampus (estimated marginal means (EMM) = 11.25 dB increase, *p_FDR_* < 0.001), middle/posterior cingulate/precuneus (EMM = 7.65 dB, *p_FDR_* = 0.013), amygdala (EMM = 6.97 dB, *p_FDR_* < 0.023). Anterior insula (EMM = 1.59 dB, *p_FDR_* = 0.73), posterior insula (EMM = 1.34 dB, *p_FDR_* = 0.73), anterior cingulate (EMM = 4.92 dB, *p_FDR_* = 0.14), and thalamus (EMM = 6.67 dB, *p_FDR_* = 0.14) did not reach significance in terms of their change at post-RC relative to baseline. It is worthwhile noting that the lack of significant changes in thalamus firing rates may be somewhat due to the limited sample size for this ROI, as the degree of increase was still relatively large. Pairwise comparisons revealed that changes in hippocampal firing rates were significantly higher than both anterior insula (*p_FDR_* < 0.001) and posterior insula (*p_FDR_* = 0.002), as well as anterior cingulate (*p_FDR_* = 0.042). The same was true for the comparison between neurons in middle/posterior cingulate/precuneus compared to posterior insula (*p_FDR_* = 0.033).

**Figure 5.**
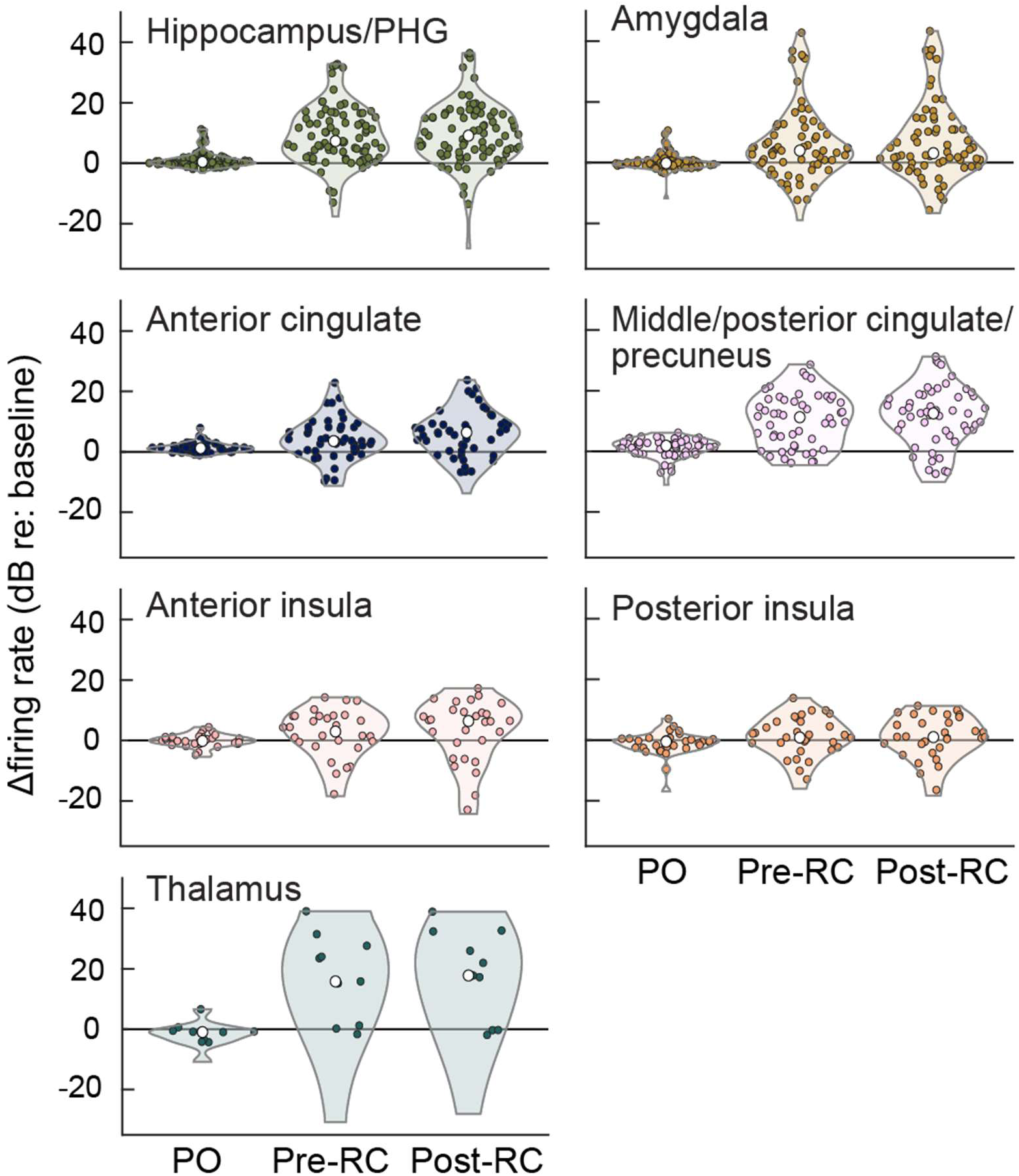
ROI-specific firing rate changes during emergence. Changes in firing rate are expressed in decibels (dB) relative to baseline firing rate (determined from a 2-minute period prior to propofol off (PO)). Each *colored symbol* represents a single neuron. *White symbols* are medians for each time window. PHG: parahippocampal gyrus. PO: propofol off. RC: response to command.

To determine the sequence of neuronal firing rate changes among the brain regions sampled, we calculated the latency to change (either increase or decrease) after cessation of propofol infusion (PO; see *Methods*). To compare across participants with different emergence times, as with the distance from criticality analysis, we again normalized time by defining *t* = 0 as PO time and *t* = 1 as RC time. This analysis was limited to those neurons that changed their firing rate during emergence, as well as those with a minimum firing rate of 0.1 Hz at all four pre-defined stages, as above (n = 240 neurons; **Figure 6; Supplementary Figure 5**). We observed that neurons in MTL were some of the earliest to change. For example, neurons in hippocampus and parahippocampal gyrus (PHG) that showed significant modulation during emergence (68 out of 81 hippocampal/PHG neurons) changed with a median *t* = 0.48 (IQR = 0.37) through emergence, with 59/68 modulated neurons showing significant changes in firing rate before RC, followed closely by amygdala with a median *t* = 0.51 (IQR = 0.27), with 42/49 modulated neurons changing before RC. The limited sampling of neurons that we had from thalamus *(n* = 11, with 9 of these significantly modulated during emergence and 8/9 modulated prior to RC) also changed early (median *t* = 0.42; IQR = 0.03). These areas were followed by middle/posterior cingulate & precuneus (median *t* = 0.64; IQR = 0.60; 34/46 modulated neurons changing prior to RC), anterior cingulate (median *t* = 0.88; IQR = 0.58; 25/33 modulated neurons changing prior to RC) and finally anterior insula and posterior insula, which on average changed with a median t = 0.90 (IQR = 0.22, 14/19 neurons) and *t* = 0.90 (IQR = 0.72, 10/16 neurons), respectively. i.e., nearly simultaneous with RC. Heterogeneous latencies were observed within ROIs (**Supplementary Figure 6**). Further, statistical analyses showed that the thalamus (EMM = 0.54 nmlz time, *p_FDR_* = 0.006), hippocampus (EMM = 0.59 nmlz time, *p_FDR_* < 0.001), amygdala (EMM = 0.68 nmlz time, *p_FDR_* = 0.004), middle/posterior cingulate and precuneus (EMM = 0.69 nmlz time, *p_FDR_* = 0.004) showed changes in firing rate significantly earlier than the time of RC. Anterior cingulate (EMM = 0.88 nmlz time, *p_FDR_* = 0.34), anterior insula (EMM = 0.82 nmlz time, *p_FDR_* = 0.23) and posterior insula (EMM = 0.97 nmlz time, *p_FDR_* = 0.84) were not significantly different from 1 (i.e., the time of RC). Pairwise comparisons on these data revealed that hippocampus (*p_FDR_* = 0.004), thalamus (*p_FDR_* = 0.029), middle/posterior cingulate/precuneus (*p_FDR_* = 0.030) and amygdala (*p_FDR_* = 0.038) showed modulation significantly earlier than posterior insula, while hippocampus was also significantly earlier than anterior cingulate (*p_FDR_* = 0.029).

**Figure 6.**
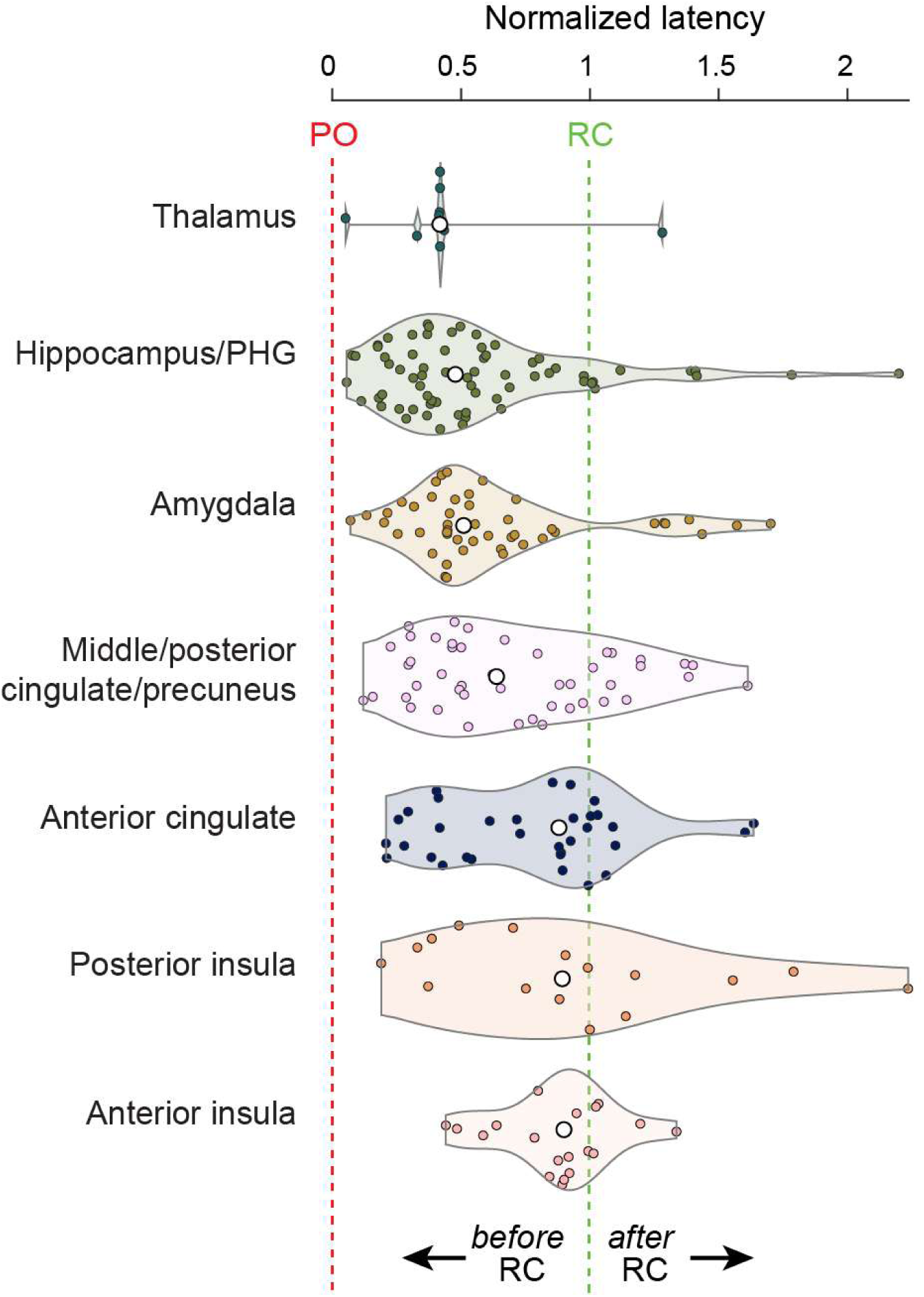
Early activation of medial temporal lobe ROIs during emergence. Latencies to change in firing rate relative to baseline for all neurons organized by ROI. Each *colored symbol* is one neuron. *White symbols* are within-ROI medians.

## Discussion

In this study, we provide the first human single-neuron characterization of the transition from propofol-induced unresponsiveness to recovery of volitional behavior. Across MTL, cingulate, and insular cortex, we observed a regional ordering in the timing of these changes, albeit with marked heterogeneity in firing rate modulation within each region. Neurons in the MTL, including hippocampus, amygdala, and PHG, showed the earliest changes from baseline (i.e., during propofol anesthesia) activity, frequently preceding the behavioral return of responsiveness. Activity changes in cingulate and insular cortices followed later and clustered nearer to the moment of behavioral recovery.

### Neural mechanisms of anesthetic emergence

Previous work on the neural mechanisms of emergence has focused on neural inertia, the hysteresis between induction and emergence that reflects active neurobiological resistance to state transitions[19, 32, 33]. Prior accounts also emphasize the contribution of subcortical arousal centers in the brainstem, hypothalamus, and thalamus[34–37] and mesocircuit interactions that regulate global cortical excitability[35, 37, 38]. Neurons in thalamus in the current study were among the earliest to change firing rates, consistent with its established role as a relay between subcortical arousal centers and cortex, though we note this observation cautiously given the limited sampling. Neuroimaging studies of emergence have provided macro-scale evidence regarding the reorganization of the brain during anesthetic emergence[39, 40]. Our data complement this literature by demonstrating that, at the level of single neurons, emergence is marked by region-specific and temporally ordered changes in firing that are not necessarily tied to behavioral responsiveness. This sequential reactivation of discrete ROIs supports the idea that emergence is an active “reboot” of brain activity[41].

The later activation of anterior insula and anterior cingulate cortex is consistent with their well-described roles in salience processing, conscious access, and facilitation of goal-directed behavior[42–44]. Prior fMRI data demonstrated association of activity in those areas with return of responsiveness following both dexmedetomidine and propofol anesthesia[39], and in the current study, the latency to changes in neural activity was just prior to return of responsiveness. By contrast, the unexpectedly early activation of MTL structures suggests their reorganization well before overt behavior returns. This pattern may reflect their tight coupling with subcortical sleep–wake circuitry[45, 46], early reinstatement of internal subjective experience in the form of dreaming, or the presence of covert consciousness when volitional motor output is still absent. Neurons in middle/posterior cingulate cortex and precuneus changed firing rates significantly earlier than RC, clustering temporally with MTL structures rather than with anterior cingulate. This earlier engagement is consistent with proposals that posterior medial cortex sustains a “core self” or baseline self-related processing that persists even at deep levels of anesthesia and is among the first to re-engage during emergence[5].

Dreaming and dream-like internal mentation are common during moderate levels of propofol anesthesia[40, 47, 48]. Furthermore, hippocampus and amygdala are consistently active during REM sleep[49, 50] and episodic memory processing during sleep[51, 52]. Indeed, anesthesia emergence has similarities with awakening from sleep. Lewis et al[53] showed that anesthesia emergence consistently includes a transient brain state characterized by K-complex-like events that are hallmarks of N2 sleep, and neural inertia has been likened to sleep inertia, the period of impaired arousal and cognition that accompanies awakening from sleep[54]. Early MTL reactivation during emergence therefore may reflect the reinstatement of internally generated subjective experience—i.e., disconnected consciousness—before the patient is able to act on commands. Indeed, prior evidence has shown that MTL regions, such as the hippocampus, are often recruited prior to reporting their conscious thoughts. For example, an fMRI study of experienced mindfulness practitioners showed increased activity occurring several seconds before spontaneous thought detection[55], with the greatest changes occurring bilaterally in hippocampus and parahippocampal gyrus, although activity was also apparent to a lesser extent in posterior cingulate, rostral anterior cingulate, inferior parietal lobule and posterior insula.

A second, not mutually exclusive possibility is that early MTL activity reflects the initial stages of reconnection with external stimuli before motor output is restored. This is consistent with a recent report indicating that hippocampus is processing sensory information even in anesthetized state[56]. Although direct evidence for covert connected consciousness during anesthesia emergence is limited, the concept has precedent in disorders of consciousness, where neuroimaging has revealed intact cognitive processing in behaviorally unresponsive patients[57, 58]. The temporal gap between MTL reactivation and that of cingulate and insular cortex, regions associated with salience processing and volitional action and previously linked to restoration of connected consciousness[40, 42], is consistent with a period in which neural processing has resumed but is not yet sufficient to drive behavioral responsiveness. Whether this intermediate state reflects disconnected consciousness, emerging connected awareness without motor capacity, or some combination remains an open question that future studies using neural decoding or post-emergence interviews could address.

### The dynamics of anesthesia emergence

Whereas the latency analysis characterizes the timing of individual neurons, we also captured the collective dynamics of the recorded population to assess emergent properties of state transitions that are not apparent from single-neuron statistics alone. We did so using two complementary approaches, measuring distance to criticality (**Figure 3**), and tracking population activity through state space (**Figure 4**).

The observed decrease in *d*_2_ during emergence provides direct evidence that the recovery of responsiveness is accompanied by a shift of neural population dynamics toward criticality. This finding is consistent with a growing body of evidence linking near-critical dynamics to the conscious waking state. Studies using detrended fluctuation analysis have shown that long-range temporal correlations in oscillatory brain activity break down during anesthesia[59, 60] and NREM sleep[61], while avalanche-based analyses have similarly demonstrated departures from power-law scaling during states of reduced consciousness [62, 63]. Reminiscent of our findings here, one study in mice found that the approach towards criticality during emergence from anesthesia occurs with a brain region-dependent time course[64].

Our data extend these findings in two important ways. First, whereas most prior work has compared steady-state epochs of wakefulness, sleep, or anesthesia, the present analysis tracks the *transition* between these states with single-neuron resolution, revealing that the approach to criticality during emergence is gradual and monotonic rather than abrupt. Second, our use of *d*2, a measure grounded in renormalization group theory and information-theoretic distance from the critical manifold[18], provides a principled quantification of proximity to criticality that goes beyond proxy measures such as power-law exponents or scaling indices, which are related to criticality but do not directly measure distance from it. Importantly, the *d*_2_ measure is reliable for short time series like our sliding window analysis here, thus accounting for the full repertoire of timescales in the population dynamics in a time resolved way. In contrast avalanche analysis typically requires long duration recordings to obtain reliable results.

By pooling population activity across neurons and examining the underlying geometric structure of the state-space trajectory, we found that neural activity followed a low-dimensional, orderly path through a restricted manifold during emergence. The trajectory exhibited two regions of high density that may correspond to attractor states, with two densely sampled regions corresponding to a propofol-unresponsiveness state and a post-return of responsiveness state. Notably, the first attractor encompassed the period both before and after propofol cessation, indicating that the neural population persisted in a suppressed configuration as propofol was beginning to clear from the brain. This observation suggests a single-neuron-level correlate of neural inertia, i.e., the network’s resistance to state transitions despite cessation of the anesthetic infusion. After a delay, the system escaped this attractor and traversed an extended, relatively low-density path through state space before settling into a second attractor following return of responsiveness. This switching between stable dynamical regimes is predicted by computational models of neural inertia and observed in animal studies of emergence[2, 19, 65, 66], where emergence from anesthesia proceeds through abrupt transitions between metastable network states rather than through a smooth return to wakefulness [67]. The pattern observed in the current study—prolonged dwell, escape, extended transit, recapture—differs somewhat from the discrete stochastic switching among several metastable states reported in this preclinical work. The discrepancy may reflect differences in species, anesthetic agent, number and distribution of recorded neurons, or genuine differences in the complexity of emergence dynamics across species and agents.

Within this dynamical framework, the marked heterogeneity of single-unit responses may reflect a stochastic, high-dimensional system reorganizing toward a new attractor[19–21]. The ordering of regional latencies and the observation that neurons in hippocampus and amygdala exerted the greatest influence on the population trajectory suggest that MTL structures disproportionately shapes the collective dynamics of emergence, with midline and insular cortices integrating into the recovered attractor only closer to behavioral responsiveness.

We note that the present analysis identifies putative attractor states qualitatively on the basis of trajectory density. Formal characterization of attractor stability, e.g., through estimation of energy landscapes or eigenvalue spectra of linearized dynamics around each state[68, 69], will require larger neuronal samples.

### Emergence from anesthesia as reassembly of the self

Sleigh and colleagues suggested that anesthesia produces a fragmentation of selfhood, in which elements of subjective experience, agency, embodiment, and memory may dissociate during induction and reassemble during emergence[70]. Their model proposes that selfhood is comprised of hierarchically organized components: a core self (the primitive feeling of existence, linked to posterior cingulate cortex), embodied self and sentience (linked to insula), agency and volition (linked to anterior cingulate and salience network), and narrative identity and autobiographical memory (linked to MTL and default mode network). During emergence, this hierarchy would be expected to reassemble from the bottom, i.e., core self and interoceptive processes first, narrative and autobiographical components last. Subcortical arousal centers, including the thalamus, would be expected to be recruited first as a necessary precondition for cortical re-engagement.

Although our behavioral measure (RC) indexes connected consciousness, not self-related processing directly, we may infer changes in self-related processing if we assume that changes in firing rate reflect specific regions previously identified with self-related processing being recruited. Under this assumption, our limited thalamic data are consistent with the latter expectation of the Sleigh et al. model. By contrast, our results are contrary to the predicted sequence of cortical reassembly in that model. Neurons in the autobiographical-memory network (hippocampus and PHG) along with amygdala reactivated earliest, preceding restoration of behavioral agency. Only later did the insula and cingulate cortex - regions tied to interoception, action selection, and volition - reactivate, marking the transition to connected consciousness and the capacity to act on commands. This raises the intriguing possibility that regions involved in autobiography and emotional processing reactivate during emergence before those involved in embodied or volitional aspects of self.

We note that early MTL firing rate changes do not necessarily imply the reinstatement of narrative selfhood. Hippocampal reactivation may reflect sleep-to-wake transitional processes such as sharp-wave ripple-associated replay[53], and early amygdala activity may be driven by ascending arousal signals rather than autobiographical or emotional processing. Distinguishing between these functional interpretations will require paradigms that directly probe self-related cognition during emergence.

The late reactivation of cingulate and insular cortex aligns with PET imaging data from Scheinin et al.[40], who identified thalamus, cingulate cortices, and angular gyri as structures fundamental for connected consciousness, independent of pharmacological agent or direction of state change. In our data, cingulate and insular neurons changed firing rates near the time of return of responsiveness, consistent with these regions gating the transition from disconnected to connected consciousness. The convergence across PET imaging in healthy volunteers and single-neuron recordings in neurosurgical patients suggests that the temporal dissociation between MTL and cingulate/insular reactivation may be a generalizable feature of anesthetic emergence.

These data raise the possibility that narrative or mnemonic aspects of self reappear before embodied or volitional aspects, implying that the internal restoration of awareness precedes its external expression. Whether or not the early MTL activity reflects narrative selfhood *per se*, the inversion of the hypothesized hierarchy highlights the value of single-unit recordings for investigating the temporal sequence of regional reactivation during emergence. This sequence would be difficult-to-impossible to capture with non-invasive neuroimaging methods (scalp EEG or fMRI) during anesthesia transitions in a clinical setting.

### Caveats and limitations

This study has several caveats and limitations. First, the sample size is modest and drawn from neurosurgical patients with epilepsy, in whom electrode placement was constrained by clinical necessity. This resulted in dense sampling of MTL, but limited coverage of regions implicated in core self-processing, such as precuneus and lateral prefrontal cortex. Consequently, whether the regional ordering we describe extends to these areas remains to be determined. Another important caveat is that our participants had medically refractory epilepsy, which may have affected the neural dynamics examined in the present study. However, cortical network changes during propofol anesthesia in this population parallel those observed during natural sleep[71], suggesting that the fundamental architecture of state transitions is preserved.

While we infer self-related processes based on known functional neuroanatomy, we did not directly assess subjective experience, covert consciousness, or self-related cognition during emergence. Finally, return of responsiveness captures volitional motor report but not the onset of subjective experience, which may have occurred earlier. Multimodal paradigms incorporating neural decoding, self-localizers, or post-emergence interviews will be required to disentangle these components.

### Future directions

Future work should explicitly compare the neural sequence of anesthetic emergence with transitions during natural sleep, where MTL activation also occurs in the absence of connected consciousness, and incorporate behavioral or decoding-based assessments of covert consciousness. The population trajectory framework developed here provides a basis for comparing emergence dynamics across anesthetic agents. Volatile agents may produce trajectories with distinct attractor geometry or regional ordering compared with propofol. Structured post-emergence interviews could test whether early MTL reactivation corresponds to dream-like disconnected experience, awareness of the external environment, or both. Finally, if MTL reactivation reliably precedes behavioral responsiveness, real-time monitoring of MTL activity could in principle provide earlier detection of returning consciousness than current behavioral or cortical EEG-based approaches.

## Acknowledgements

We are grateful to Haiming Chen, Christopher Garcia, Matthew Howard, Ariane Rhone, Anna Schwarting, Marek Solvik, Cameron Brace, Beril Mat, Mariel Kalkach Aparicio, Aaron Suminski, and Dillon Scott for help with data collection, analysis, interpretation, and helpful comments on the manuscript.

## Author contributions

Conceptualization: JIB, MIB, KVN

Methodology: JIB, RNM, CAA Software: JIB, WLS

Formal analysis: JIB

Investigation: JIB, MIB, ERD, UJG, RNM, HK, CAA, MB, WLS, KVN

Data Curation: JIB, ERD, KVN

Visualization: JIB, MIB, KVN

Writing - Original Draft: JIB, MIB, KVN

Writing - Review & Editing: JIB, MIB, ERD, UJG, RNM, HK, CAA, MB, WLS, KVN

Funding acquisition: MIB, MB, KVN

## Data, code, and materials availability

Files and code to reproduce the main figures in this manuscript will be available at https://doi.org/10.5281/zenodo.20146616. Due to confidentiality, raw patient data are not made publicly available. Further anonymized data may be available on request dependent on data transfer agreements and other required collaborative documents. Code for *d*_2_ calculation is publicly available at https://github.com/sam-sooter/prox_crit_toolkit.

## Supplementary Materials

### Materials and Methods

#### Data collection

##### Ethics Statement

Research protocols were approved by the University of Iowa and University of Wisconsin Institutional Review Boards (protocol #201804807 “Consciousness Research”). Written informed consent was obtained from all participants. Research participation did not interfere with acquisition of clinically necessary data, and participants could rescind consent for research without interrupting their clinical management.

##### Participants and experimental procedure

This study included 11 adult neurosurgical patients (8 female; ages 21-61 years old, median age 44 years old) implanted with electrodes in temporal, parietal, and frontal cortex to identify epileptic foci (**Supplementary Table 1**). Patients were studied between February 2023 and July 2025 at The University of Iowa Hospitals and Clinics and at the University of Wisconsin Hospital and Clinics.

Extracellular single neuron recordings were obtained in the operating room just after the electrode implantation surgery during emergence from propofol anesthesia. Induction of general anesthesia was accomplished with propofol, fentanyl and rocuronium. Surgical level of anesthesia was initially maintained with sevoflurane, followed by propofol and remifentanil with the goal to achieve emergence solely from propofol at the end. Electrode implantation was performed under robotic assistance (ROSA One Brain robotic platform, Zimmer Biomet, Warsaw, IN). Following implantation of the last electrode array, muscle relaxant was reversed, remifentanil administration was stopped, and electrodes were connected to the data acquisition hardware (see below). Once electrophysiology data collection commenced, propofol infusion continued for five minutes at a rate of 50-200 µg/kg/min and then turned off (“propofol off”, PO). Anesthesiologists were instructed to cease narcotics (e.g. fentanyl, remifentanil, hydromorphone) and anxiolytics prior to neural recordings and to not give these again until recordings were completed. Following PO, patients were monitored continuously for spontaneous movement and assessed for response to verbal commands at approximately 1-minute intervals by a research team member. Return of responsiveness was defined as the first response to command (RC) in a sequence of two consecutive positive responses. Recordings continued for a minimum of 5 minutes, as clinical circumstances permitted.

Because this was an observational study and these recordings were opportunistic, patients were neither consecutive in time nor selected randomly. Inclusion in the dataset was determined by the availability of single neuron data recorded at specific time points relative to surgery. Patients were included when recordings were continuous from five minutes prior to propofol cessation to at least two minutes following return of responsiveness, and where single neurons were able to be resolved offline. The single neuron data presented here are from 11 participants who met these inclusion criteria.

##### Electrodes and recording system

Stereo electroencephalography (sEEG) depth electrode arrays (Ad-Tech Medical, Oak Creek, WI) were placed in brain locations based solely on a clinical need to identify seizure foci[72]. For the purpose of recording single neurons, sEEG electrode arrays were of a hybrid design (i.e., ‘Behnke-Fried’ electrodes)[73]. These hybrid arrays included eight high impedance microwires (39 µm diameter; platinum-iridium) that were insulated plus one uninsulated microwire. The microwires were cut to extend beyond the end of the macro sEEG probe by 2 to 4 mm, depending on the distance of the most distal macro contact to the appropriate brain target.

Each of these microwires was splayed individually immediately prior to implantation. Electrode locations were confirmed based on post-operative MRI scans, preprocessed using FreeSurfer [74] (see below for further details). Up to eight hybrid electrodes were implanted in each patient. Neurophysiological data were recorded using a Neuralynx Atlas System (Neuralynx, Bozeman, MT). High impedance recordings were first passed through a preamplifier located on top of the patient’s head (ATLAS-HS-36-CHET-A9, Neuralynx, Bozeman, MT) prior to interfacing with the ATLAS acquisition system. Data were recorded at a 32000 Hz sampling rate, filtered between 0.1 – 8000 Hz and referenced online to the uninsulated microwire.

##### Imaging

A T1-weighted structural MRI scan of the brain was conducted for each participant both before and after the implantation of electrodes. The images were captured using a 3T Siemens TIM Trio scanner and a 12-channel head coil. MPRAGE images had a spatial resolution of 0.78 × 0.78 mm, a slice thickness of 1.0 mm, and utilized a repetition time of 2.53 s and an echo time of 3.52 ms. To locate the recording positions on the preoperative structural MRI scans, these images were aligned with post-implantation structural MRIs. This alignment was achieved using a 3D linear registration algorithm (Functional MRI of the Brain Linear Image Registration Tool[75]) and custom-written MATLAB scripts (MathWorks, Natick, MA). Included microwire bundles were verified to be within gray matter. For visualization purposes only, recording site locations were co-registered for each participant to a template brain to derive MNI coordinates. Coordinates of microwire locations were then visualized on an fsaverage MRI template using custom-written MATLAB scripts.

#### Data analysis

##### Single neuron analysis

Data from high impedance electrodes were denoised using the demodulated band transform[76], downsampled to 12 kHz, and common average re-referenced to all high impedance contacts on the same microwire bundle prior to spike sorting. Spike sorting was performed using an automated procedure with manual curation, utilizing an algorithm implementing high-order spectral decomposition to aid in pattern recognition and identify individual features [28, 77]. Briefly, filters were estimated for candidate single neuron waveforms for each channel through a process related to blind deconvolution. Extracted features were clustered using a Gaussian mixture model in MATLAB (R2022a, MathWorks Inc) and spike times from these clustered features were used to identify candidate waveforms.

Single neurons were then manually curated and defined based on classical waveform shapes, consistency of waveforms across time for each cluster, and interspike interval distributions displaying refractory periods (<1% of interspike intervals occurring within 1 ms). Putative single neuron spike times were then referenced to the PO time. For each neuron, spike counts were binned across time into 1 s windows, converted to firing rates (Hz) and then smoothed with a 5 s moving average window across the whole recording. Transient outliers were removed based on values >2 standard deviations above mean firing rates within a 100 s moving window and replaced with this mean value. Firing rates were *z*-scored with respect to a baseline period of 120 s immediately prior to propofol cessation. The latency of change relative to baseline firing following propofol cessation was determined based on the time at which a neuron’s firing rate showed an absolute z-score >2.5, with an additional condition that this threshold was also surpassed between 20 to 60 s after this time period, in order to further avoid periods of transient increases in firing rate in determining this timing.

To examine data across participants, where time to recovery was variable across individuals, spike data were normalized by calculating the difference between RC time and PO time (“duration to emergence”) and then reformatting the corresponding emergence time vector related to the binned spike data (in 1 second bins) for each participant by subtracting the PO time and dividing by this duration to emergence. In this scenario, zero represented the time of PO and 1 represented the first of the two consecutive RCs, allowing for any timepoints prior to PO to be <0 and any following RC to be >1. Binned firing rates (1 s bins) for each participant were then interpolated onto a common time vector that ranged from −0.3 to 1.3 (allowing for up to 30% of the duration to emergence prior to PO and following RC). Data from participants that did not have 30% of the duration covered before and after were substituted with NaNs and the final common normalized time period covered across individuals ranged from −0.1330 to 1.1505. All neurons were *z*-score normalized across this entire window prior to performing PCA analyses (utilizing the “pca” function in Matlab). Density in PCA space (see **Figure 4b**) was calculated by computing a two-dimensional histogram of the corresponding state-space data.

##### Distance to criticality analysis

ROIs were included in the distance to criticality analysis where for each participant a minimum of five single neurons were isolated. Spike times for each neuron were first binned with a 0.2 s resolution and corresponding spike counts were concatenated across neurons for a particular ROI. *d*_2_ (i.e. distance to criticality, where 0 = precisely at criticality) was estimated utilizing the method provided by[18]. Neuronal data were analyzed in 40-second non-overlapping sliding windows. The autoregressive model order was set at 16. *d*_2_ values were normalized based on the method described above prior to grouping data across participants (see **Figure 3**).

##### Statistical analysis

To examine differences in firing rates and response latencies across different time periods and ROIs, linear mixed effects (LME) models were fit to the corresponding values, using the *fitlme* function in Matlab with restricted maximum likelihood estimation enabled. For firing rate changes, only neurons with a minimum firing rate of 0.1 Hz at each time period of interest were included. The LME model for firing rate ratios (relative to baseline) included a fixed effect of ROI and a random effect of participant, in the form:

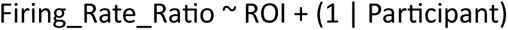

Estimated marginal means from this model for each ROI were tested for significance against zero via the *coeftest* function to determine whether changes in firing rates were significantly elevated overall relative to pre-PO values. For the latency analysis, neurons were only included that increased their firing rates based on the criteria described above during the course of emergence. The LME model for latency changes again included a fixed effect of ROI and a random effect of participant, in the form:

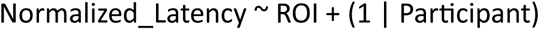

Estimated marginal means from the normalized latency model were tested against 1 (i.e., RC) to determine whether changes across neurons were significantly earlier than RC for each ROI. Pairwise comparisons between ROIs for both models were performed based on the *coefTest* function implemented with different contrasts applied, so that weights of +1 and −1 were assigned to each ROI pair comparison. Corresponding *p*-values for each model (for both the reference tests and the contrast tests) were corrected for multiple comparisons with a Benjamini-Hochberg false discovery rate procedure.

---

**Supplementary Table 1.** Participant demographics and clinical information. Other drugs given during the OR procedure are specified b [Add other drugs as column].

| Participant | Age | Sex | Antiseizure medications | Seizure focus and notes | *n*<sub>units</sub> |
| --- | --- | --- | --- | --- | --- |
| Pt1 | 36 | F | cenobamate, lamotrigine, levetiracetam XR, topiramate XR | R medial temporal | 32 |
| Pt2 | 59 | F | gabapentin, levetiracetam, oxcarbazepine | R medial temporal; prior L temporal lobectomy | 49 |
| Pt3 | 36 | M | cannabidiol, clobazam, lacosamide, levetiracetam | Multifocal with unclear onset; prior R anterior temporal lobectomy | 18 |
| Pt4 | 44 | F | cenobamate, zonisamide | L medial temporal | 31 |
| Pt5 | 48 | F | lamotrigine, levetiracetam, lorazepam | L medial temporal | 23 |
| Pt6 | 21 | M | cenobamate, clobazam, lacosamide, oxcarbazepine, zonisamide | B multifocal with unclear onset | 21 |
| Pt7 | 46 | M | alprazolam, cenobamate, lamotrigine, lorazepam, zonisamide | R inferior frontal and L medial temporal | 24 |
| Pt8 | 61 | F | brivaracetam, cenobamate, pregabalin | B medial temporal | 79 |
| Pt9 | 56 | F | clobazam, clonazepam<br>eslicarbazepine,<br>gabapentin,<br>levetiracetam | R frontal; R frontal<br>cortical dysplasia | 40 |
| Pt10 | 32 | F | clobazam,<br>oxcarbazepine,<br>zonisamide | L cingulate | 18 |
| Pt11 | 36 | F | cenobamate,<br>lamotrigine, clobazam | L posterior<br>heterotopia | 21 |

**Supplementary Figure 1.**
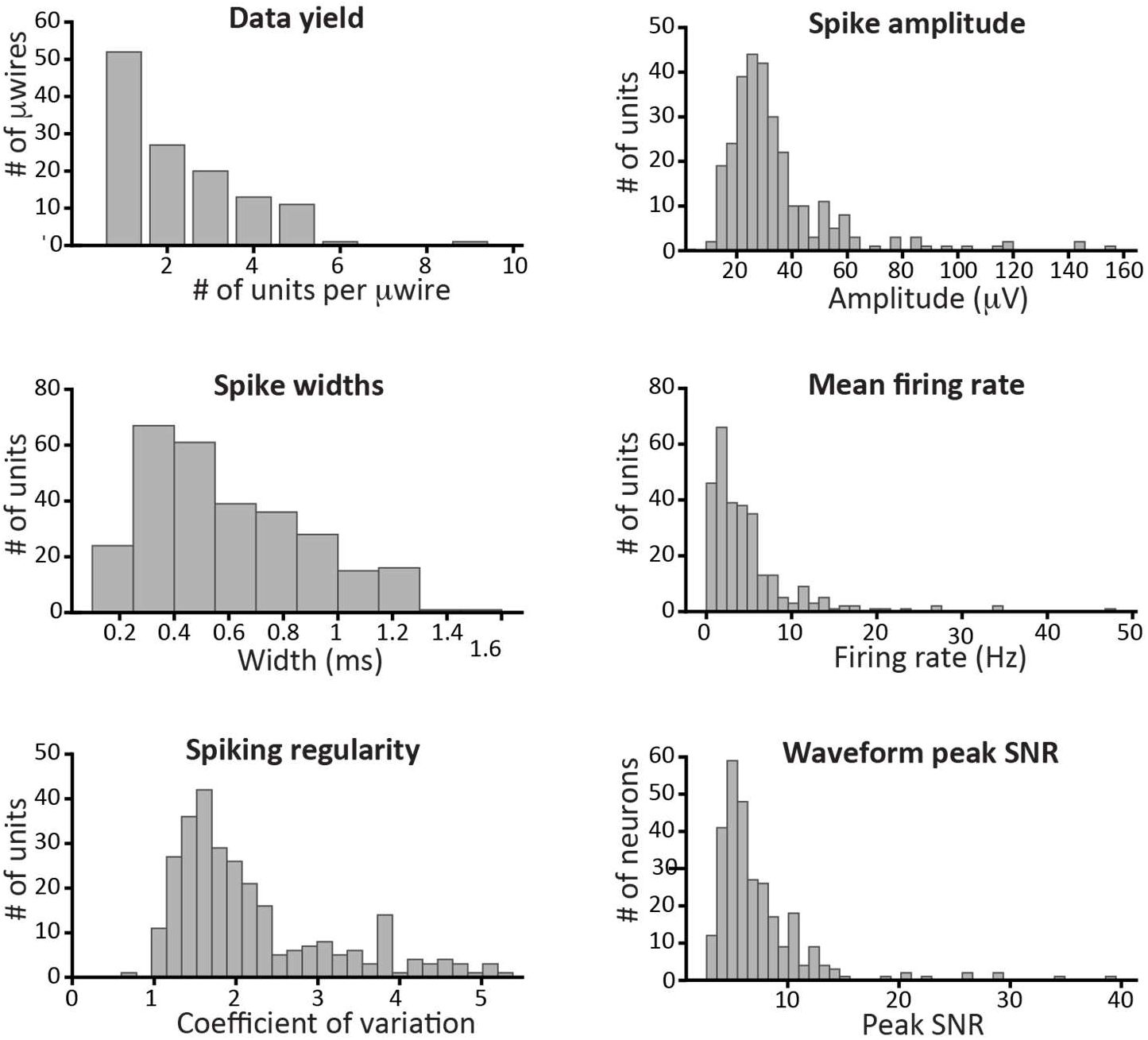
Single neuron metrics. Shown are histograms of data yield per microwire, spike amplitude, spike width, mean firing rate, spiking regularity, and waveform peak signal-to-noise ratio (SNR).

**Supplementary Figure 2.**
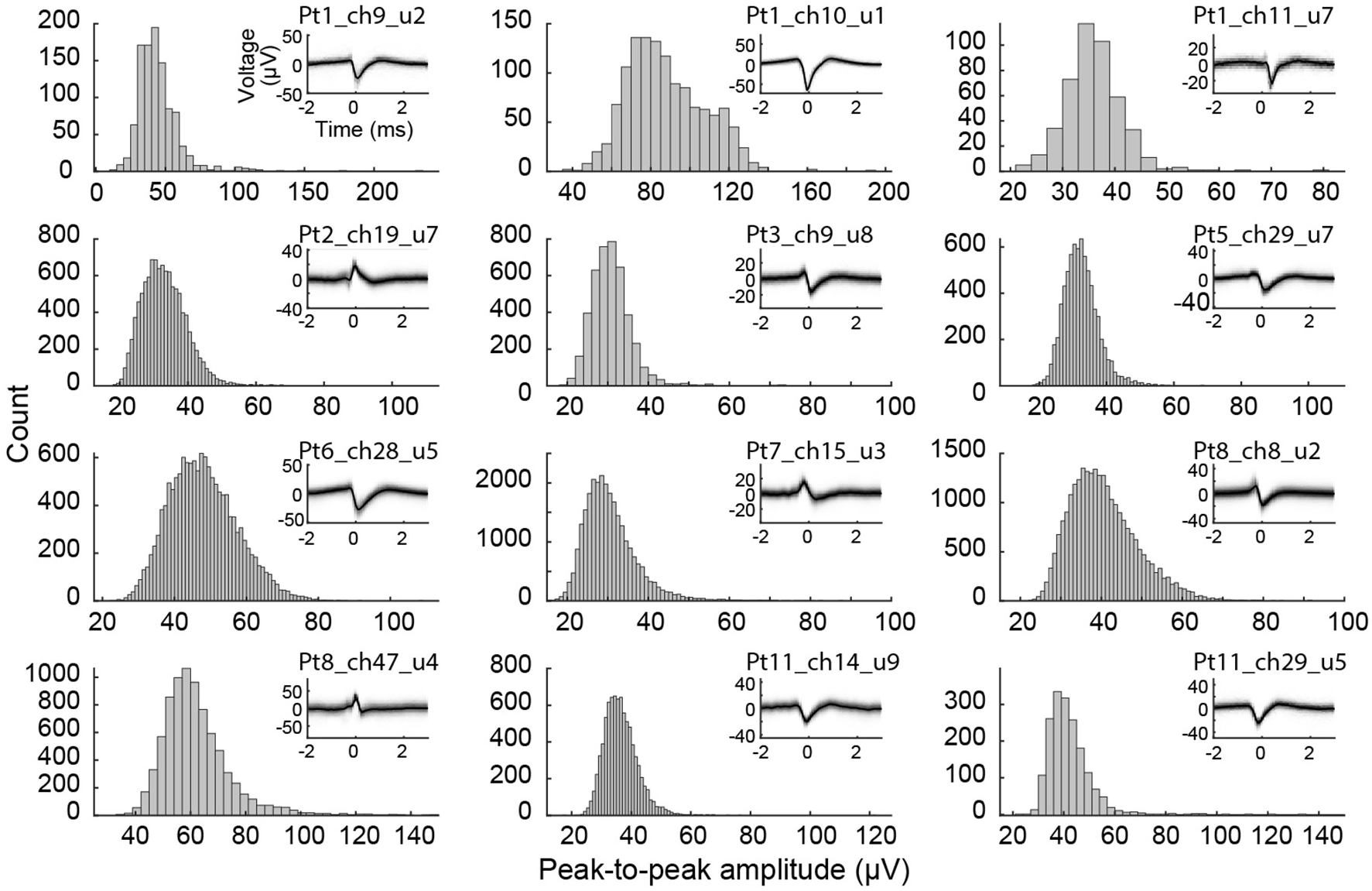
Single neuron spike amplitude distributions. Amplitude distributions (*histograms*) and unit waveforms (*insets*) are shown for twelve randomly selected isolated neurons.

**Supplementary Figure 3.**
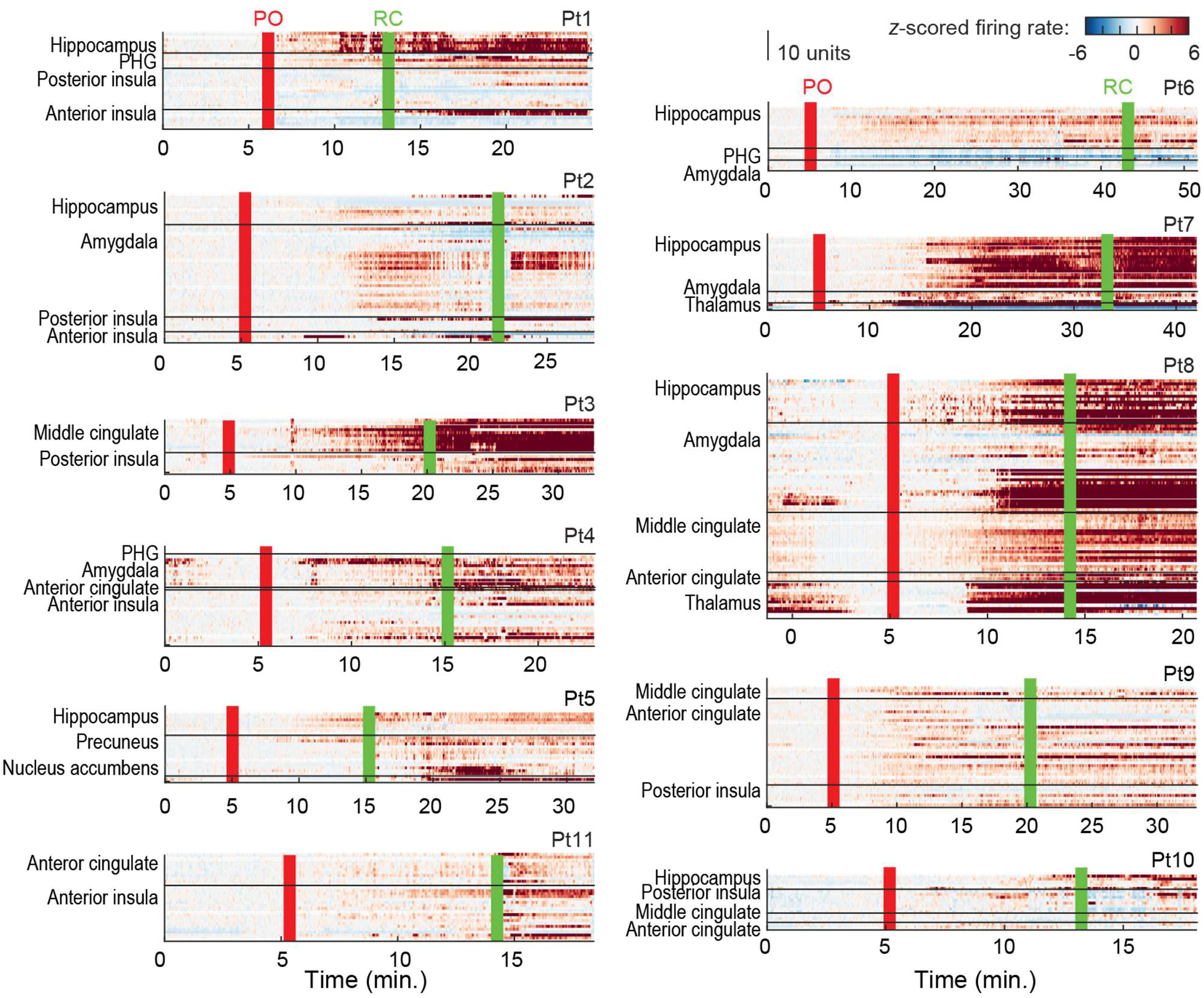
Firing rate data for all neurons in the dataset. Firing rate for all neurons recorded in each participant. Each row corresponds to one neuron. Firing rates were *z*-scored and are shown as heat maps.

**Supplementary Figure 4.**
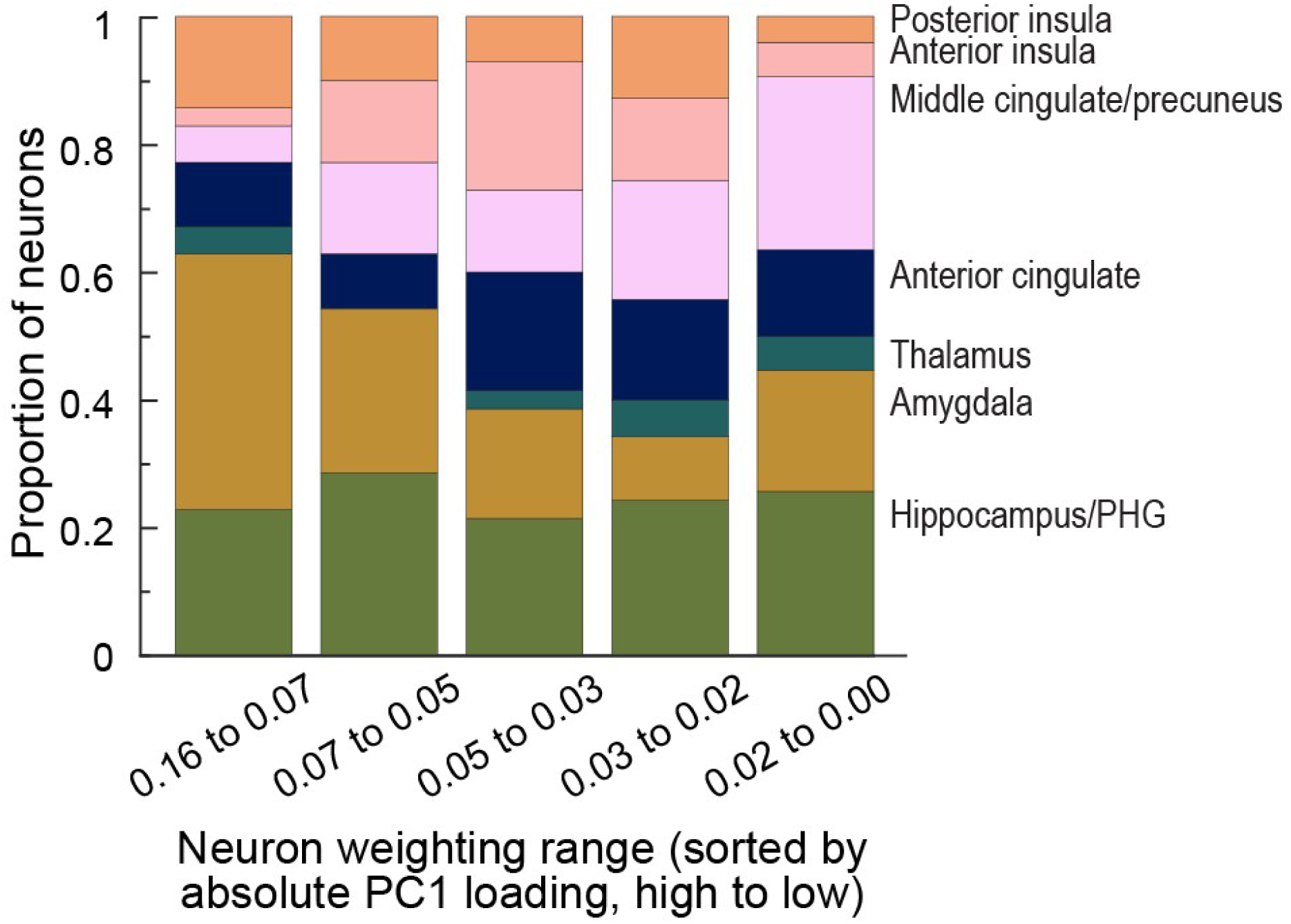
PCA loadings by region. Contributions of neurons in each region to the first principal component of the emergence trajectory (PC1).

**Supplementary Figure 5.**
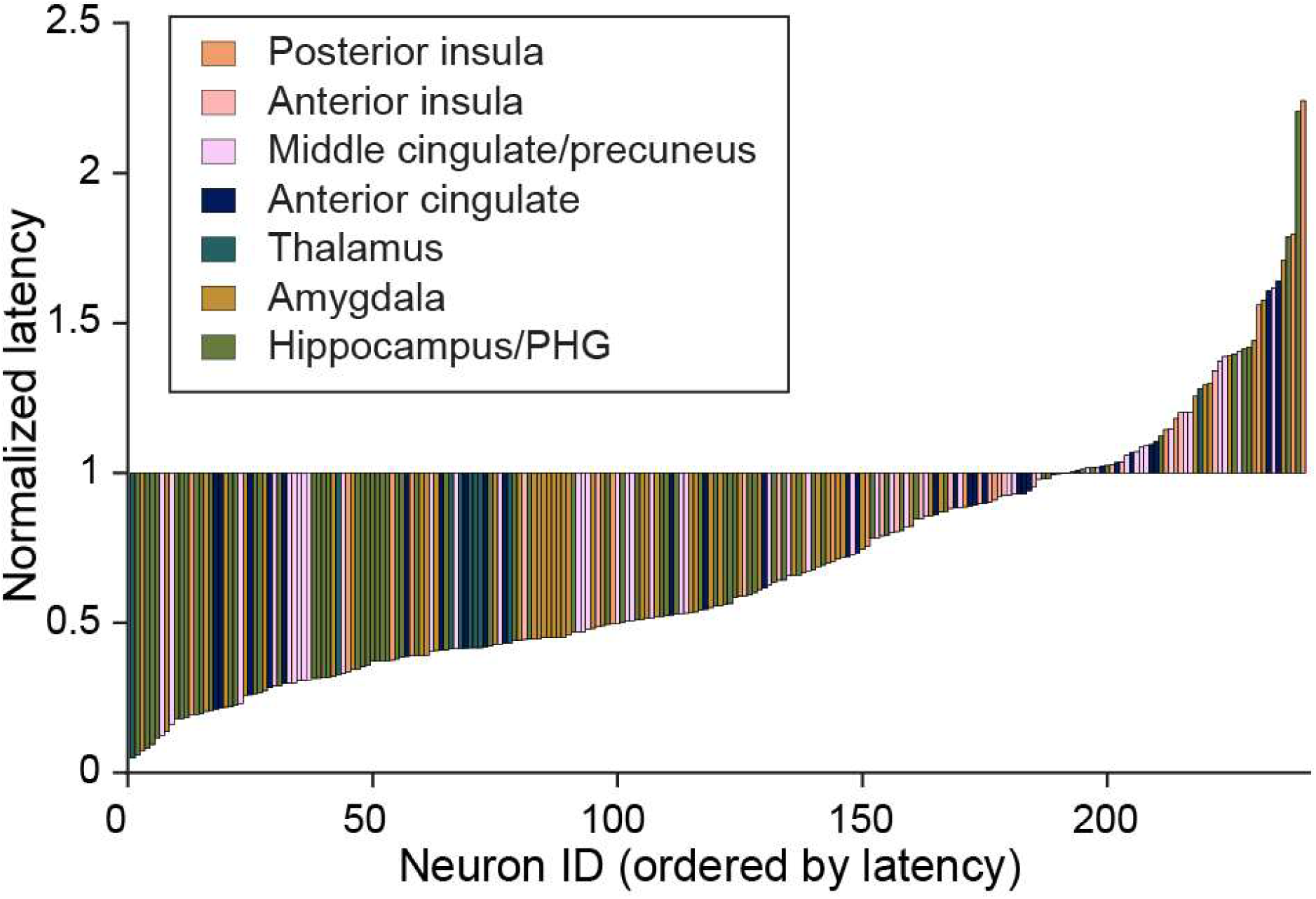
Normalized latency of change in firing rate for all single neurons. Each bar corresponds to one neuron, with location coded by color (see *inset*).

**Supplementary Figure 6.**
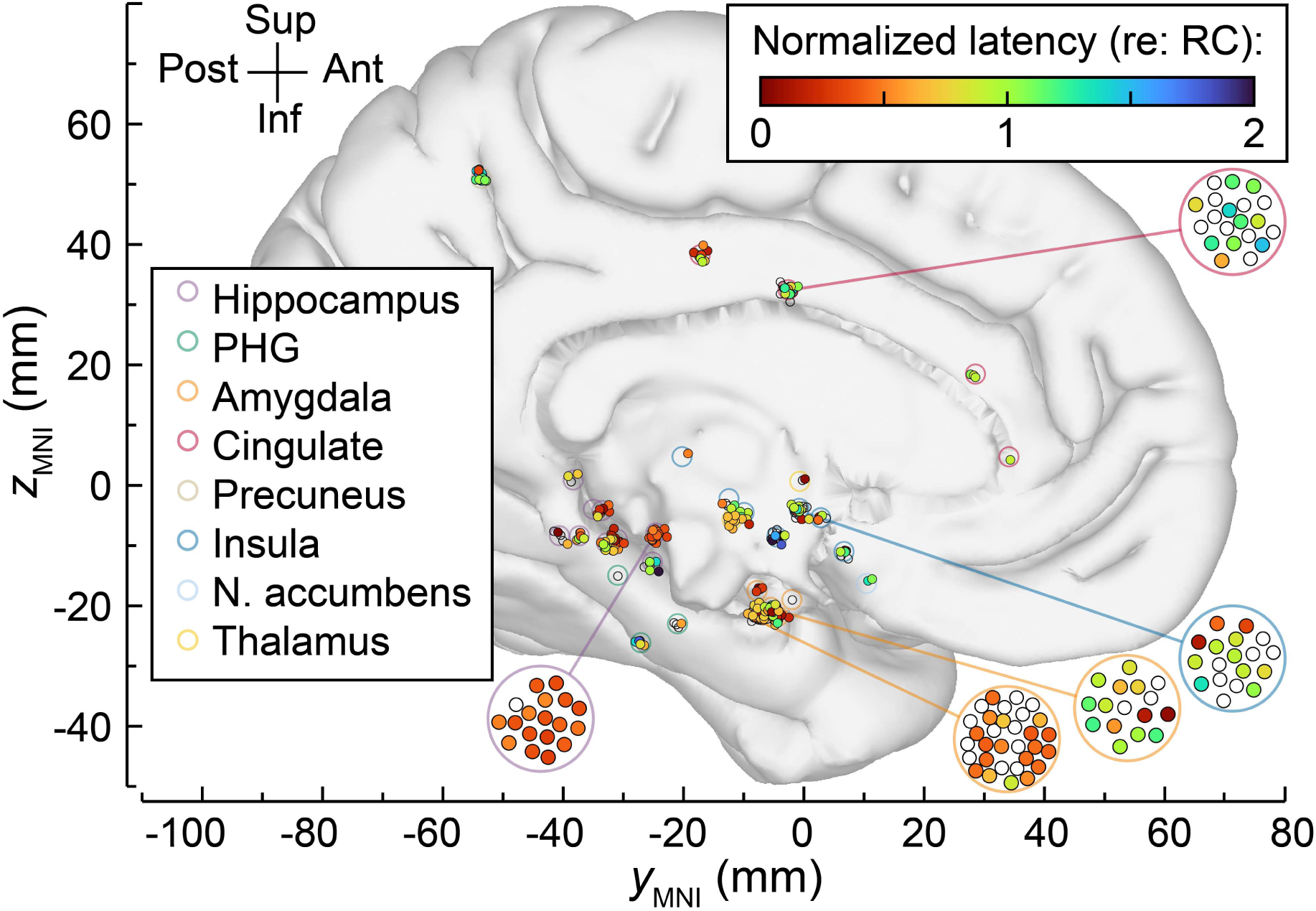
Anatomical distribution of latencies during emergence. Each *small filled circle* corresponds to one neuron, with the color indicating the normalized latency. *Large unfilled circles* mark the locations of microwire bundles in the common MNI space, with colors indicating brain region.

